# Regulation of Plant Phototropic Growth by NPH3/RPT2-like Substrate Phosphorylation and 14-3-3 Binding

**DOI:** 10.1101/2021.04.09.439135

**Authors:** Stuart Sullivan, Thomas Waksman, Louise Henderson, Dimitra Paliogianni, Melanie Lütkemeyer, Noriyuki Suetsugu, John M. Christie

## Abstract

Polarity underlies all plant physiology and directional growth responses such as phototropism. Yet, our understanding of how plant tropic responses are established is far from complete. The plasma-membrane associated BTB-containing protein, NON-PHOTOTROPIC HYPOCOTYL 3 (NPH3) is a key determinant of phototropic growth which is regulated by AGC kinases known as the phototropins (phots). However, the mechanism by which phots initiate phototropic signalling via NPH3, and other NPH3/RPT2-like (NRL) members, has remained unresolved. Here we demonstrate that NPH3 is directly phosphorylated by phot1 both *in vitro* and *in vivo*. Light-dependent phosphorylation within a conserved consensus sequence (RxS) located at the extreme C-terminus of NPH3 is necessary to promote its functionality for phototropism and petiole positioning in *Arabidopsis*. Phosphorylation of this region by phot1 also triggers 14-3-3 binding combined with changes in NPH3 phosphorylation and localisation status. Seedlings expressing mutants of NPH3 that are unable to bind or constitutively bind 14-3-3s show compromised functionality that is consistent with a model where signalling outputs arising from a gradient in NPH3 RxS phosphorylation/localisation across the stem are a major contributor to phototropic responsiveness. Our current findings provide further evidence that 14-3-3 proteins are instrumental components regulating auxin-dependent growth and show for the first time that NRL proteins are direct phosphorylation targets for plant AGC kinases. Moreover, the C-terminal phosphorylation site/14-3-3-binding motif of NPH3 is conserved in several members of the NRL family, suggesting a common mechanism of regulation.

## Introduction

The ability to sense and respond to the prevailing light conditions is instrumental for plants to adapt their growth and development to the external environment. Phototropism allows plants to re-orientate shoot growth towards a directional light source, which promotes light capture and early seedling growth (Christie and Murphy, 2013). Phototropism is induced by UV/blue light and is mediated by two phototropin (phot) receptor kinases, phot1 and phot2 (Fankhauser and Christie, 2015). Phot1 is the primary phototropic receptor and functions over a wide range of fluence rates, whereas phot2 activity requires higher light intensities (Sakai et al., 2001). Phots also control physiological responses such as chloroplast movement, leaf positioning, leaf expansion and stomatal opening (Christie, 2007), which together serve to optimize photosynthetic efficiency and growth (Takemiya et al., 2005; Gotoh et al., 2018; Hart et al., 2019).

Phototropins are plasma membrane-associated kinases containing two light, oxygen, or voltage-sensing domains (LOV1 and LOV2) at their N-terminus, which bind oxidized flavin mononucleotide (FMN) as a UV/blue light absorbing cofactor (Christie et al., 1999; Sullivan et al., 2008). Light perception, primarily by LOV2, results in activation of phototropin kinase activity and receptor autophosphorylation (Christie et al., 2002; Cho et al., 2007). Although multiple phosphorylation sites have been identified within phot1 and phot2 (Christie et al., 2015), sites within the kinase activation loop are important for signalling, and kinase-inactive variants of phot1 and phot2 are non-functional (Inoue et al., 2008b; Inoue et al., 2011). Despite the importance of phot kinase activity for downstream signalling, only a limited number of substrates have been identified to date. BLUE LIGHT SIGNALING 1 (BLUS1) and CONVERGENCE OF BLUE LIGHT AND CO_2_ 1 (CBC1) are phot1 kinase substrates involved in blue-light induced stomatal opening (Takemiya et al., 2013; Hiyama et al., 2017), while phosphorylation of ATP-BINDING CASSETTE B19 (ABCB19) and PHYTOCHROME KINASE SUBSTRATE 4 (PKS4) by phot1 modulates hypocotyl phototropism (Christie et al., 2011; Demarsy et al., 2012; Schumacher et al., 2018). Given the variety of physiological responses mediated by phot signalling, further phot kinase substrates likely await identification (Schnabel et al., 2018).

Phototropism results from the establishment of lateral gradients of the phytohormone auxin, which leads to increased cell expansion on the shaded side of the hypocotyl (Christie and Murphy, 2013). NON-PHOTOTROPIC HYPOCOTYL 3 (NPH3) is an essential signalling component for phototropism and is required for the formation of the lateral auxin gradients (Motchoulski and Liscum, 1999; Haga et al., 2005). NPH3, together with ROOT PHOTOTROPISM 2 (RPT2), are the founding members of the NPH3/RPT2-Like (NRL) protein family, which contains 33 members in *Arabidopsis* (Pedmale et al., 2010; Christie et al., 2018). The primary amino acid structure of NPH3 can be separated into three regions based on sequence conservation with other NRL proteins: an N-terminal BTB (bric-a-brac, tramtrack, and broad complex) domain, a central NPH3 domain and a C-terminal coiled-coil domain (Christie et al., 2018). The C-terminal portion of NPH3, including the coiled-coil domain, is proposed to facilitate localisation of NPH3 to the plasma membrane (Inoue et al., 2008a) as well as mediating direct interaction with phot1 (Motchoulski and Liscum, 1999). NPH3 is reported to function as a substrate adapter in a CULLIN3-based E3 ubiquitin ligase complex targeting phot1 for ubiquitination (Roberts et al., 2011). Ubiquitination of phot1 may be involved in receptor desensitisation, particularly under high light irradiation (Roberts et al., 2011), but its importance in phot1 signalling is currently unknown.

Although the biochemical function of NPH3 remains unresolved, activation of phot1 by blue light results in dynamic changes to NPH3 phosphorylation status and subcellular localisation (Haga et al., 2015; Sullivan et al., 2019). NPH3 is phosphorylated on multiple sites in darkness, including sites located towards the N-terminus (Tsuchida-Mayama et al., 2008), and localises to the plasma membrane (Haga et al., 2015). Upon blue light perception, NPH3 is rapidly dephosphorylated (Pedmale and Liscum, 2007) and becomes internalised into aggregates, which transiently attenuates its interaction with phot1 (Haga et al., 2015; Sullivan et al., 2019). These effects are reversible in darkness, with the kinetics of NPH3 rephosphorylation matching the photoactive lifetime of phot1 (Hart et al., 2019). The kinases and phosphatases which modulate NPH3 phosphorylation status are unknown, however reduced levels of dephosphorylation, and relocalisation into aggregates, correlates with enhanced phototropic responsiveness observed in de-etiolated (green) seedlings (Sullivan et al., 2019).

Along with NPH3, two other NRL family members also have known roles in phot signalling pathways. RPT2 interacts with both phot1 and NPH3 (Inada et al., 2004; Sullivan et al., 2009), it is proposed to influence NPH3 phosphorylation status and promote the reconstitution of the phot1-NPH3 complex to sustain signalling under higher light intensities (Haga et al., 2015). In line with this, phototropic responsiveness in mutant seedlings lacking RPT2 decreases as light intensity is increased (Sakai et al., 2000). Similarly, *RPT2* expression levels are low in darkness, but increase with irradiation in a fluence-dependent manner (Sakai et al., 2000). RPT2, together with NPH3, is also involved in phot-mediated leaf positioning and leaf expansion responses (Inoue et al., 2008a; Harada et al., 2013). NRL PROTEIN FOR CHLOROPLAST MOVEMENT 1 (NCH1) is positioned within the same clade as RPT2 in the *Arabidopsis* NRL phylogenetic tree (Christie et al., 2018). NCH1 and RPT2 redundantly mediate chloroplast accumulation movements in response to low intensity light (Suetsugu et al., 2016).

Phot signalling is dependent upon reversible changes in phosphorylation (Christie et al., 2015). 14-3-3 proteins are present in all eukaryotic organisms and bind to target proteins through identification of phospho-serine/threonine motifs (Aitken et al., 1992; Johnson et al., 2010). 14-3-3 binding can produce a variety of consequences, such as regulation of enzymatic activity, changes in subcellular localisation, protein stability or alteration of protein-protein interactions (Camoni et al., 2018). 14-3-3 proteins are known to bind to phot1 and phot2 following receptor autophosphorylation (Kinoshita et al., 2003; Inoue et al., 2008b; Sullivan et al., 2009; Tseng et al., 2012), while NPH3 and RPT2 have both been identified as components of the 14-3-3 interactome (Schoonheim et al., 2007; Keicher et al., 2017). However, the functional relevance of these interactions and the roles of 14-3-3 proteins in phot signalling remain unclear.

Despite the importance of NRL proteins in blue-light mediated responses, how signalling is initiated upon phot activation is still not known. In the present study we identify NPH3 as a substrate for phot1 kinase activity. Phosphorylation of NPH3 at the C-terminus by phot1 results in 14-3-3 binding, which is required for early signalling events and promotes NPH3 functionality. The C-terminal phosphorylation site of NPH3 is conserved in several NRL family members, including RPT2, suggesting phot-mediated phosphorylation and 14-3-3 binding may represent a conserved mechanism of regulation.

## Results

### Light-dependent 14-3-3 binding to NPH3

In order to identify additional components involved in blue light signalling, GFP-NPH3 was immunoprecipitated from etiolated *nph3* mutant seedlings expressing functional *NPH3::GFP-NPH3* (Sullivan et al., 2019). Anti-GFP immunoprecipitations (IPs) were performed on total protein extracts from seedlings maintained in darkness or after a brief blue light treatment (20 μmol m^−2^ s^−1^ for 15 min) to capture early signalling events. Co-purifying proteins were analysed by label-free quantitative tandem mass spectrometry (MS) to allow identification of proteins whose abundance changed following blue light irradiation. As expected, phot1 was recovered in the immunoprecipitations from both dark- and light-treated seedlings, but at a higher abundance in the dark (Table S1). This is in agreement with previous results showing NPH3-phot1 interactions are attenuated by blue light (Haga et al., 2015). Conversely, several 14-3-3 isoforms were detected at greater abundance following blue light irradiation (Fig. 1A).

**Fig. 1.**
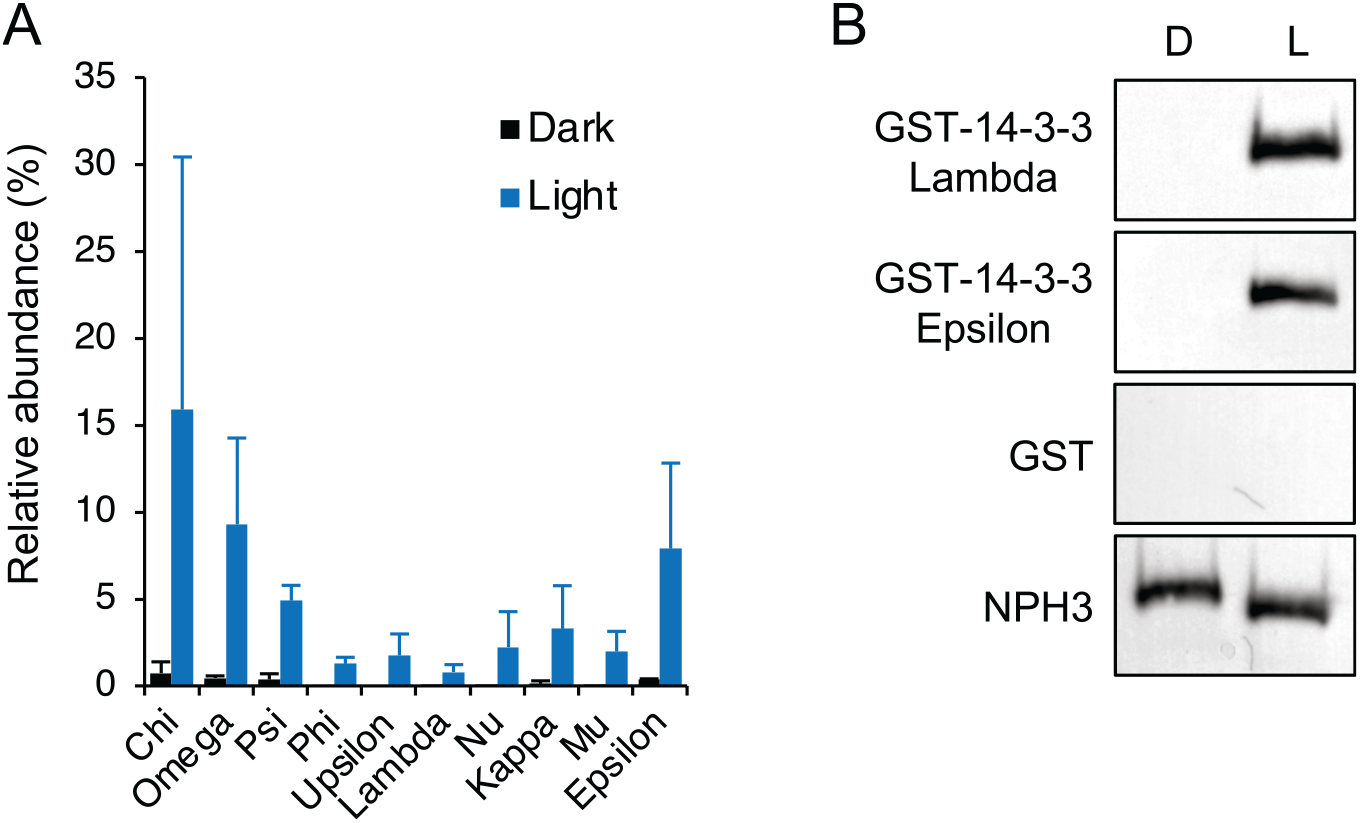
NPH3 interacts with 14-3-3 proteins in a light-dependent manner. (A) NPH3 interacting proteins were identified by mass spectrometry analysis of anti-GFP immunoprecipitations from etiolated seedlings expressing GFP-NPH3 maintained in darkness (Dark) or irradiated with 20 μmol m^−2^ s^−1^ of blue light for 15 min (Light). Each value is the mean and S.D. from two biological replicates. Protein signal intensities were converted to relative abundance of the bait protein (GFP-NPH3). (B) Far-western blot analysis of anti-GFP immunoprecipitations from etiolated seedlings expressing GFP-NPH3 maintained in darkness (D) or irradiated with 20 μmol m^−2^ s^−1^ of blue light for 15 (L). GST-tagged 14-3-3 isoforms (Lambda and Epsilon) or GST alone were used as probes. Blots were probed with anti-GFP antibody as loading control (bottom panel).

14-3-3 proteins bind to target proteins through recognition of phospho-serine/threonine containing motifs. *Arabidopsis* expresses 13 different 14-3-3 isoforms which can be phylogenetically divided into the epsilon and non-epsilon groups (DeLille et al., 2001). Far-western blotting was performed to assess direct 14-3-3 binding to GFP-NPH3. Binding of recombinant 14-3-3 Lambda (non-epsilon group member) and 14-3-3 Epsilon (epsilon group member) fused to glutathione-S-transferase (GST) was not detected for GFP-NPH3 IPs from etiolated seedlings maintained in darkness (Fig. 1B). Blue-light irradiation results in an enhanced electrophoretic mobility of GFP-NPH3 due to its rapid dephosphorylation (Pedmale and Liscum, 2007). Concurrently, binding of 14-3-3 Lambda and Epsilon was observed following irradiation, while no binding was observed when GST alone was used as the probe. In line with the results from IP-MS analysis, no specificity in binding of 14-3-3 proteins from epsilon and non-epsilon groups was detected. These results suggest that blue light irradiation triggers both phosphorylation, and concomitant 14-3-3 binding, as well as dephosphorylation events on NPH3.

### Analysis of phosphorylation sites within NPH3

Activation of phot1 by blue light results not only in rapid changes in the phosphorylation status of NPH3 but also its subcellular localisation (Haga et al., 2015; Sullivan et al., 2019). In darkness, NPH3 localises predominantly to the plasma membrane but is rapidly internalised into aggregates upon blue-light treatment. Based on data from global phosphoproteomics experiments (Durek et al., 2010; Willems et al., 2019) three regions of NPH3 (M1, M2 and M3) containing the majority of experimentally identified phosphopeptides were selected for mutational analysis (Fig. 2A). Within each of the regions, all of the serine and threonine residues were replaced with alanine to mimic the dephosphorylated state. The mutations were introduced into the *NPH3::GFP-NPH3* construct, transiently expressed in the leaves of *Nicotiana benthamiana* and compared with the expression of the non-mutated GFP-NPH3 control. Transfected *N. benthamiana* plants were dark-adapted before confocal observation. The localisation of transiently expressed GFP-NPH3 was similar to that of functionally active GFP-NPH3 in *Arabidopsis* (Sullivan et al., 2019) described above, and repeated scanning with the 488-nm laser used to concomitantly excite GFP along with endogenous phot1 induced relocalisation of GFP-NPH3 into aggregates (Fig. 2B). The localisation of each of the transiently expressed NPH3 phospho-mutants was the same as GFP-NPH3 when imaged immediately (scan 1). Repeated laser scanning was effective in inducing relocalisation for both M1 and M2 constructs, whereas the M3 mutant failed to show any light-induced changes in subcellular localisation.

**Fig. 2.**
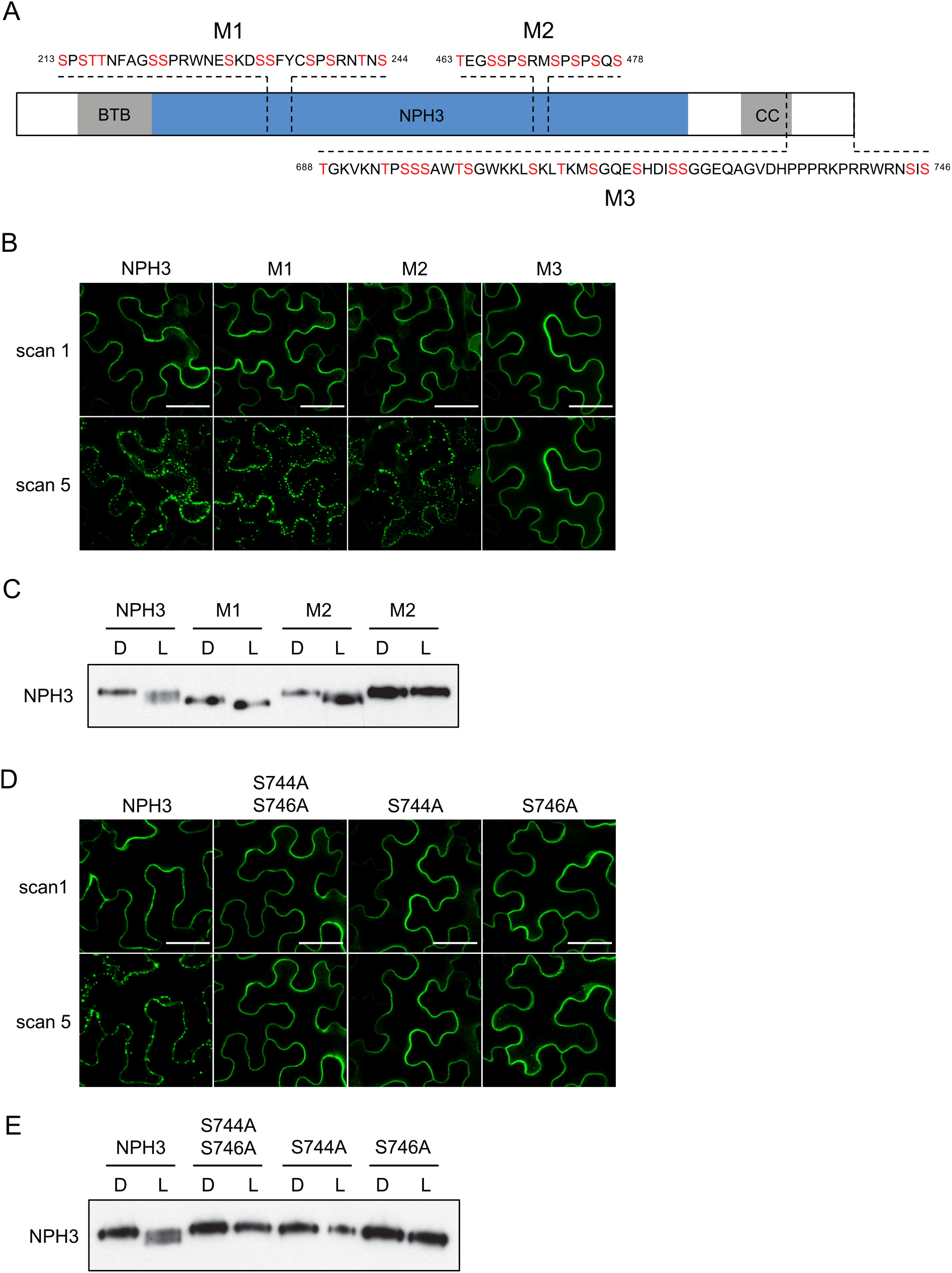
Mutational analysis of NPH3 phosphorylation sites. (A) Schematic illustration of the NPH3 protein indicating the location of the three mutagenized regions (M1, M2 and M3). For each region, all serine and threonine residues in red were substituted with alanine. The relative positions of the bric-a-brac, tramtrack, and broad complex (BTB), NPH3 and coiled-coil (CC) domains are indicated. (B) Confocal images of GFP-NPH3 (NPH3) and phosphorylation site mutants (M1, M2 and M3) transiently expressed in leaves of *N. benthamiana*. Plants were dark-adapted before confocal observation and images acquired immediately (scan 1) and after repeat scanning with the 488 nm laser (scan 5). Bar, 50 μm. (C) Immunoblot analysis of protein extracts from leaves of *N. benthamiana* transiently expressing GFP-NPH3 (NPH3) and phosphorylation site mutants (M1, M2 and M3). Plants were dark-adapted and maintained in darkness (D) or irradiated with 20 μmol m^−2^ s^−1^ of blue light for 15 min (L). Protein extracts were probed with anti-GFP antibodies. (D) Confocal images of GFP-NPH3 (NPH3) and phosphorylation site mutants S744A S746A, S744A and S746A transiently expressed in leaves of *N. benthamiana*. Plants were dark-adapted before confocal observation and images acquired immediately (scan 1) and after repeat scanning with the 488 nm laser (scan 5). Bar, 50 μm. (E) Immunoblot analysis of protein extracts from leaves of *N. benthamiana* transiently expressing GFP-NPH3 (NPH3) and phosphorylation site mutants S744A S746A, S744A and S746A. Plants were dark-adapted and maintained in darkness (D) or irradiated with 20 μmol m^−2^ s^−1^ of blue light for 15 min (L). Protein extracts were probed with anti-GFP antibodies.

Phot1-induced changes in NPH3 localisation are correlated with changes in NPH3 phosphorylation status in transgenic *Arabidopsis* seedlings (Haga et al., 2015; Sullivan et al., 2019). Immunoblot analysis of protein extracts from dark-adapted leaves of *N. benthamiana* transiently expressing GFP-NPH3 irradiated with blue light also showed an enhanced electrophoretic mobility compared to leaves maintained in darkness (Fig. 2C), although to a lesser degree than observed in etiolated *Arabidopsis* seedlings expressing GFP-NPH3 when equivalent light treatments were used (Fig. 1B). Both the M1 and M3 mutants were affected for this response, whereas the M2 mutant response was similar to GFP-NPH3 (Fig. 2C). The M1 mutant showed enhanced electrophoretic mobility in the dark compared to the GFP-NPH3 construct, with a further slight enhancement following blue light treatment. The M1 mutant contains mutations of serine residues S213, S223, S233 and S237, mutation of which was previously shown to contribute to reducing the electrophoretic mobility of NPH3 in darkness (Tsuchida-Mayama et al., 2008). Conversely, the M3 mutant migrated at the same position as GFP-NPH3 in the dark, even following blue light irradiation. Therefore, amino acid residues within the M3 region at the C-terminus of NPH3 are required for both relocalisation and dephosphorylation in response to blue light.

The C-terminal amino acid sequence of NPH3 is highly conserved in angiosperms (Fig. S1A) and contains two serine residues, S744 and S746 in *Arabidopsis* NPH3. Mutation of either serine residue to alanine, singularly or together, prevented (for S744A and S744A S746A) or greatly reduced (for S746A) the light-induced relocalisation response when transiently expressed in *N. benthamiana* (Fig. 2D). Similarly, these mutations also prevented dephosphorylation of NPH3 following blue light irradiation (Fig. 2E). Therefore, mutation of S744 and/or S746 can reproduce the results obtained with the M3 mutant. While serine to alanine mutations effectively block phosphorylation of the respective residue, phosphomimetic substitutions aim to mimic the phosphorylated state by replacement with a negatively charged amino acid. However, mutation of S744 and S746 to aspartate produced similar results to the alanine mutations; loss of light-induced relocalisation and dephosphorylation (Fig. S1B, Fig. S1C).

### S744 is required for 14-3-3 binding and early signalling events

To examine the effects of the C-terminal serine residues S744 and S746 in NPH3 signalling, we generated transgenic *Arabidopsis* expressing *NPH3::GFP-NPH3* containing S744A S746A, S744D S746D, S744A or S746A mutations in the *nph3* mutant background. Confocal imaging of hypocotyl cells of etiolated seedlings expressing S744A S746A or S744D S746D showed that both mutants did not relocalise into aggregates following irradiation with the 488nm laser, in contrast to the GFP-NPH3 control (Fig. 3A). The single S744A mutant also lacked this response, whereas the S746A mutant was unaffected. Furthermore, analysis of NPH3 dephosphorylation showed that seedlings expressing S744A S746A, S744D S746D or S744A exhibited no change in electrophoretic mobility with blue light treatment, in contrast to S746A and GFP-NPH3 expressing lines, which both displayed an enhanced mobility with blue light treatment (Fig. 3B). Whereas results from transient expression analysis in *N. benthamiana* showed both S744 and S746 were involved in these early signalling responses (Fig. 2D, Fig. 2E), analysis of transgenic *Arabidopsis* identifies only S744 as being required.

**Fig. 3.**
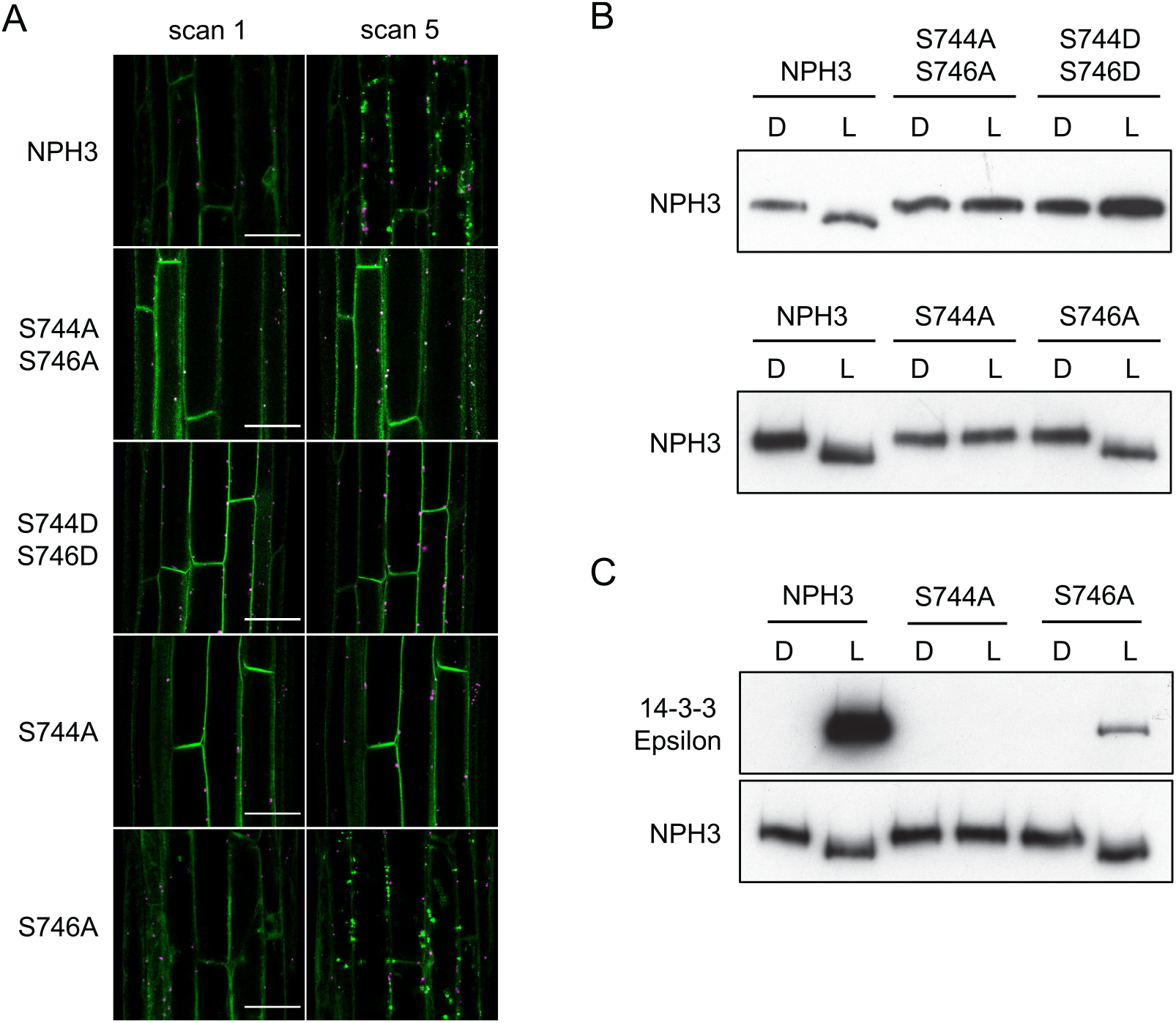
S744 is required for 14-3-3 binding and early signalling events. (A) Confocal images of hypocotyl cells of etiolated seedlings expressing GFP-NPH3 (NPH3) or phosphorylation site mutants S744A S746A, S744D S746D, S744A and S746A. Seedlings were scanned immediately (scan 1) and again after repeat scanning with the 488 nm laser (scan 5). GFP is shown in green and autofluorescence in magenta. Bar, 50 μm. (B) Immunoblot analysis of total protein extracts from etiolated seedlings expressing GFP-NPH3 (NPH3) or phosphorylation site mutants maintained in darkness (D) or irradiated with 20 μmol m^−2^ s^−1^ blue light for 15 min (L). Protein extracts were probed with anti-NPH3 antibodies. (C) Far-western blot analysis of anti-GFP immunoprecipitations from etiolated seedlings expressing GFP-NPH3 (NPH3) or phosphorylation site mutants (S744A or S746A) maintained in darkness (D) or irradiated with 20 μmol m^−2^ s^−1^ of blue light for 15 min blue light (L). GST-tagged 14-3-3 isoform Epsilon was used as the probe. Blots were probed with anti-NPH3 antibody as loading control (bottom panel).

To determine whether S744 was also required to mediate interactions between NPH3 and 14-3-3 proteins, far-western blotting was performed on anti-GFP IPs from seedlings expressing GFP-NPH3 or GFP-NPH3 containing S744A or S746A mutations (Fig. 3C). Binding of recombinant 14-3-3 Epsilon was evident for both GFP-NPH3 and S746A in a light-dependent manner, with the signal for S746A being substantially lower. However, no binding could be detected for the S744A mutant. Phosphorylation of S744 is therefore necessary for 14-3-3 binding, subcellular relocalisation and dephosphorylation of N-terminal sites (including S213, S223, S233 and S237) in response to blue light perception.

### Phot1 phosphorylates NPH3 at position S744 in a light-dependent manner

Given the evidence for light-induced phosphorylation of NPH3, we examined whether NPH3 was a direct substrate for phot1 kinase activity using a gate-keeper engineered phot1 (phot1^GK^), that can accommodate the bulky ATP analogue N^6^-benzyl-ATPγS as a thiophospho-donor (Schnabel et al., 2018). NPH3, or the NPH3 S744A mutant, were co-expressed in a cell-free expression system with phot1^GK^ and used for *in vitro* kinase assays in the presence of N^6^-benzyl-ATPγS. Light-induced thiophosphorylation, which can be detected by immunoblotting with anti-thiophosphoester antibody (α-TPE) following chemical alkylation of the incorporated thiophosphates, was detected for NPH3 but not for the S744A mutant (Fig. 4A), showing phot1 can specifically phosphorylate residue S744 of NPH3 *in vitro*. To detect the phosphorylation status of S744 *in vivo* we raised a phospho-specific antibody (pS744). Phosphorylation of S744 was observed in WT and GFP-NPH3 expressing seedlings in a light-dependent manner and mutation of S774 resulted in a loss of signal demonstrating the specificity of the pS744 phospho-specific antibody (Fig. 4B). Phosphorylation of S744 was also detectable for S746A expressing seedlings at a reduced level, similar to the results observed for 14-3-3 binding (Fig. 3C).

**Fig. 4.**
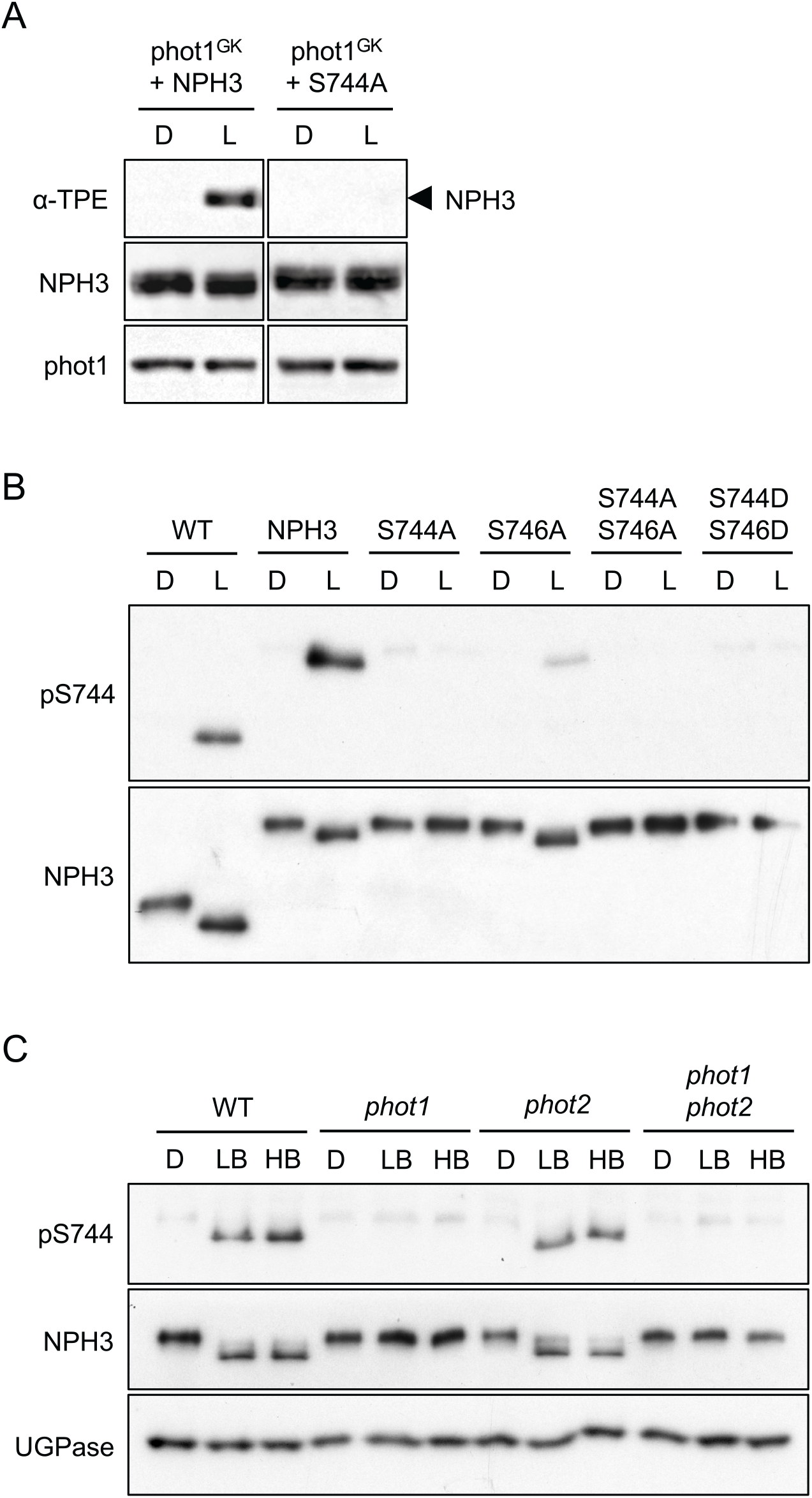
Phot1 phosphorylates NPH3 at position S744 in a light-dependent manner. (A) Thiophosphorylation analysis of *in vitro* kinase assays containing gatekeeper engineered phot1 (phot1^GK^) and NPH3 or NPH3-S744A. Reactions were performed in the absence (D) or presence of 20 s of white light (L), and thiophosphorylation was detected using anti-thiophosphoester antibody (α-TPE). Blots were probed with anti-NPH3 and anti-phot1 antibodies. (B) Immunoblot analysis of total protein extracts from etiolated seedlings expressing GFP-NPH3 (NPH3) or phosphorylation site mutants maintained in darkness (D) or irradiated with 20 μmol m^−2^ s^−1^ blue light for 15 min (L). Protein extracts were probed with phospho-specific pS744 antibody and anti-NPH3 antibodies. (C) Immunoblot analysis of total protein extracts from etiolated wild-type (WT) or *phot1, phot2* and *phot1 phot2* mutant seedlings maintained in darkness (D) or irradiated with 0.5 μmol m^−2^ s^−1^ (low blue; LB) or 50 μmol m^−2^ s^−1^ (high blue; HB) of blue light for 60 min. Blots were probed with phospho-specific pS744 antibody, anti-NPH3 anti-UGPase (loading control) antibodies.

Phot1 is the main photoreceptor mediating phototropism to low (<1 μmol m^−2^ s^−1^) and high (>1 μmol m^−2^ s^−1^) fluence rates of blue light, whereas phot2 functions predominantly at higher light intensities (>10 μmol m^−2^ s^−^1; Sakai et al., 2001). Phosphorylation of S744 occurred in WT seedlings in response to both low blue (0.5 μmol m^−2^ s^−1^) and high blue (50 μmol m^−2^ s^−1^) light treatments concomitantly with NPH3 dephosphorylation, detected via changes in electrophoretic mobility when probed with anti-NPH3 antibody (Fig. 4C). These responses were absent in *phot1 phot2* double mutant and *phot1* single mutant seedlings, but unchanged in the *phot2* single mutant, demonstrating that phosphorylation of S744 and dephosphorylation of NPH3 are phot1-specific responses in etiolated seedlings.

To assess the kinetics of changes in NPH3 phosphorylation status we performed time-course experiments. Complete dephosphorylation of NPH3 required 15 min of blue light irradiation (Fig. 5A), whereas phosphorylation of S744 was detected within 30 s and maintained over the 2 h irradiation period. When etiolated seedlings were returned to darkness following blue light exposure, S744 was dephosphorylated within 15 min, matching the time required for rephosphorylation of sites responsible for the electrophoretic mobility shift (Fig. 5B). Therefore, phot1 phosphorylation of S744 is rapid, occurring before light-induced dephosphorylation, and reversible in darkness.

**Fig. 5.**
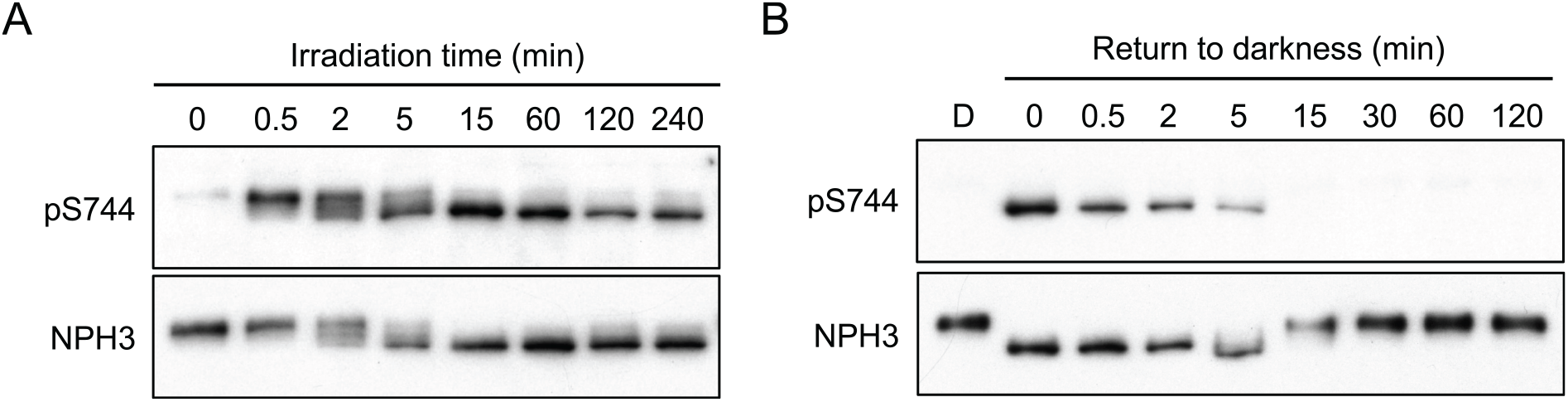
Kinetics of phot1-mediated phosphorylation of NPH3. (A) Time-course of S744 phosphorylation. Immunoblot analysis of total protein extracts from etiolated wild-type seedlings irradiated with 0.5 μmol m^−2^ s^−1^ of blue light for the time indicated. Blots were probed with anti-pS744 and anti-NPH3 antibodies. (B) Time-course of S744 dephosphorylation. Immunoblot analysis of total protein extracts from etiolated wildtype seedlings maintained in darkness (D) or irradiated with 0.5 μmol m^−2^ s^−1^ for 15 min and returned to darkness for the time indicated. Blots were probed with anti-pS744 and anti-NPH3 antibodies.

### Phot1 phosphorylation of NPH3 promotes functionality

*Arabidopsis* mutants lacking NPH3 fail to exhibit hypocotyl phototropism under a variety of different light conditions (Liscum and Briggs, 1996; Sakai et al., 2000). Phototropism in two independent homozygous transgenic *nph3* mutants expressing *NPH3::GFP-NPH3* is restored to levels comparable to non-transgenic WT seedlings when irradiated with 0.5 μmol m^−2^ s^−1^ of unilateral blue light (Fig. 6A). In contrast, the magnitude and kinetics of phototropic curvature was reduced in seedlings expressing GFP-NPH3 with both S744 and S746 residues mutated to alanine or aspartate (Fig. 6A). Similarly, phototropism was reduced in seedlings expressing GFP-NPH3 containing the S744A mutant, while the S746A expressing seedlings were fully functional (Fig. 6B). To determine whether the reduced phototropic responsiveness of the S744A mutant is due to altered photosensitivity, phototropism was further assessed under lower (0.05 μmol m^−2^ s^−1^; Fig. 6C) and higher (20 μmol m^−2^ s^−1^; Fig. 6D) intensity blue light irradiation. Under both fluence rates, transgenic lines expressing the S744A mutant were less responsive than the GFP-NPH3 or S746A expressing lines.

**Fig. 6.**
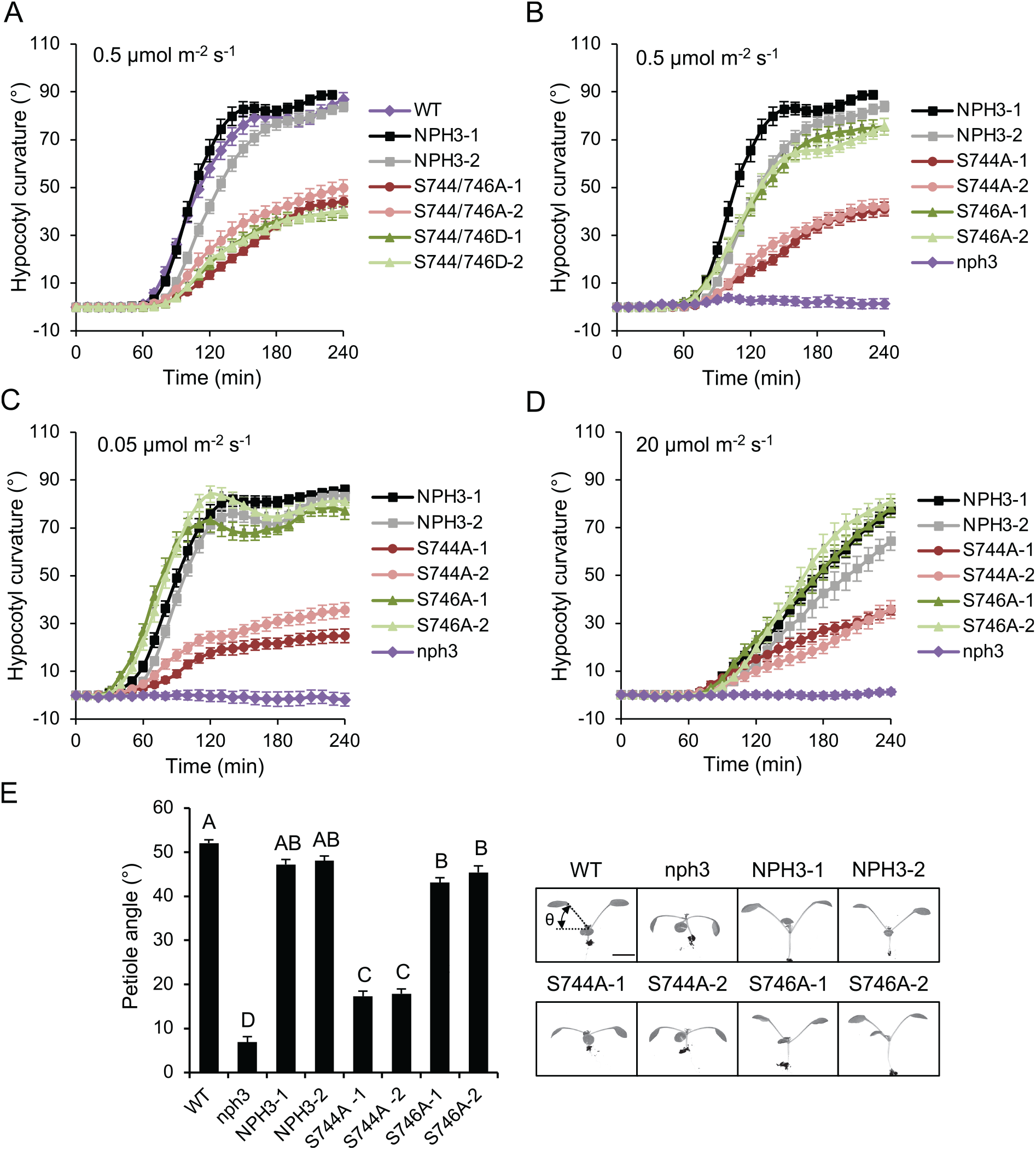
Phot1 phosphorylation of NPH3 promotes functionality. (A) Phototropism of etiolated wild-type (WT) seedlings, seedlings expressing GFP-NPH3 (NPH3) or phosphorylation site mutants S744A S746A and S744D S746D irradiated with 0.5 μmol m^−2^ s^−1^ unilateral blue light. (B – D) Phototropism of etiolated seedlings expressing GFP-NPH3 (NPH3), phosphorylation site mutants S744A or S746A and *nph3* mutant seedling irradiated with (B) 0.5 μmol m^−2^ s^−1^, (C) 0.05 μmol m^−2^ s^−1^ or (D) 20 μmol m^−2^ s^−1^ unilateral blue light. Hypocotyl curvatures were measured every 10 min for 4 h, and each value is the mean ± SE of 17-20 seedlings. (E) Petiole positioning of WT, *nph3* mutant, and seedlings expressing GFP-NPH3 or phosphorylation site mutants S744A and S746A. Plants were grown under 80 μmol m^−2^ s^−1^ white light for 9 d before transfer to 10 μmol m^−2^ s^−1^ white light for 5 d. Petiole angle from the horizontal was measured for the first true leaves, each value is the mean ± SE of 20 seedlings. Means that do not share a letter are significantly different (P < 0.01, one-way ANOVA with Tukey HSD post-test). Representative images for each genotype are shown on the right. Bar, 5 mm.

NPH3 also functions in phototropin-mediated leaf positioning, particularly in low light environments (Inoue et al., 2008a). In WT seedlings transferred to low intensity white light (10 μmol m^−2^ s^−1^) the petioles of the first true leaves were positioned obliquely upwards in order to maximise light capture, while the petioles of *nph3* mutant seedlings were positioned horizontally (Fig. 6E). Seedlings expressing GFP-NPH3 or the S746A mutant were complemented for petiole positioning, while the response of seedlings expressing the S744A mutant was significantly reduced (Fig. 6E), which was also observed for the S744A S746A and S744D S746D transgenic lines (Fig. S2). These results demonstrate that phot1 phosphorylation of S744 positively regulates NPH3 function.

### Phosphorylation and 14-3-3 binding drives NPH3 relocalisation

The phenotypes of seedlings expressing GFP-NPH3 containing S744D S746D mutations were identical to seedlings expressing NPH3 with the S744A S746A mutations (Fig. 3, Fig. 6A), consistent with reports that aspartate does not effectively mimic phosphorylation with respect to 14-3-3 binding (Maudoux et al., 2000; Johnson et al., 2010). To create a constitutively 14-3-3 binding variant, the sequence encoding the last three amino acids of the *NPH3::GFP-NPH3* construct, including the S744 phosphorylation site, was replaced with the R18 peptide sequence (Fig. 7A). R18 is a synthetic peptide that mediates phosphorylation-independent binding of 14-3-3 proteins with high affinity (Wang et al., 1999). As a control, a construct containing a mutated version of the R18 sequence (mR18), known to abolish 14-3-3 binding (Ramm et al., 2006), was also generated (Fig. 7A). When transiently expressed in *N. benthamiana*, GFP-NPH3-R18 appeared as aggregates when dark-adapted leaves were imaged immediately, with no change in localisation during imaging (Fig. 7B). Conversely, GFP-NPH3-mR18 remained localised to the plasma membrane following repeated laser scanning, as previously observed with GFP-NPH3 constructs lacking the S744 phosphorylation site (Fig. 2D, Fig. S1B).

**Fig. 7.**
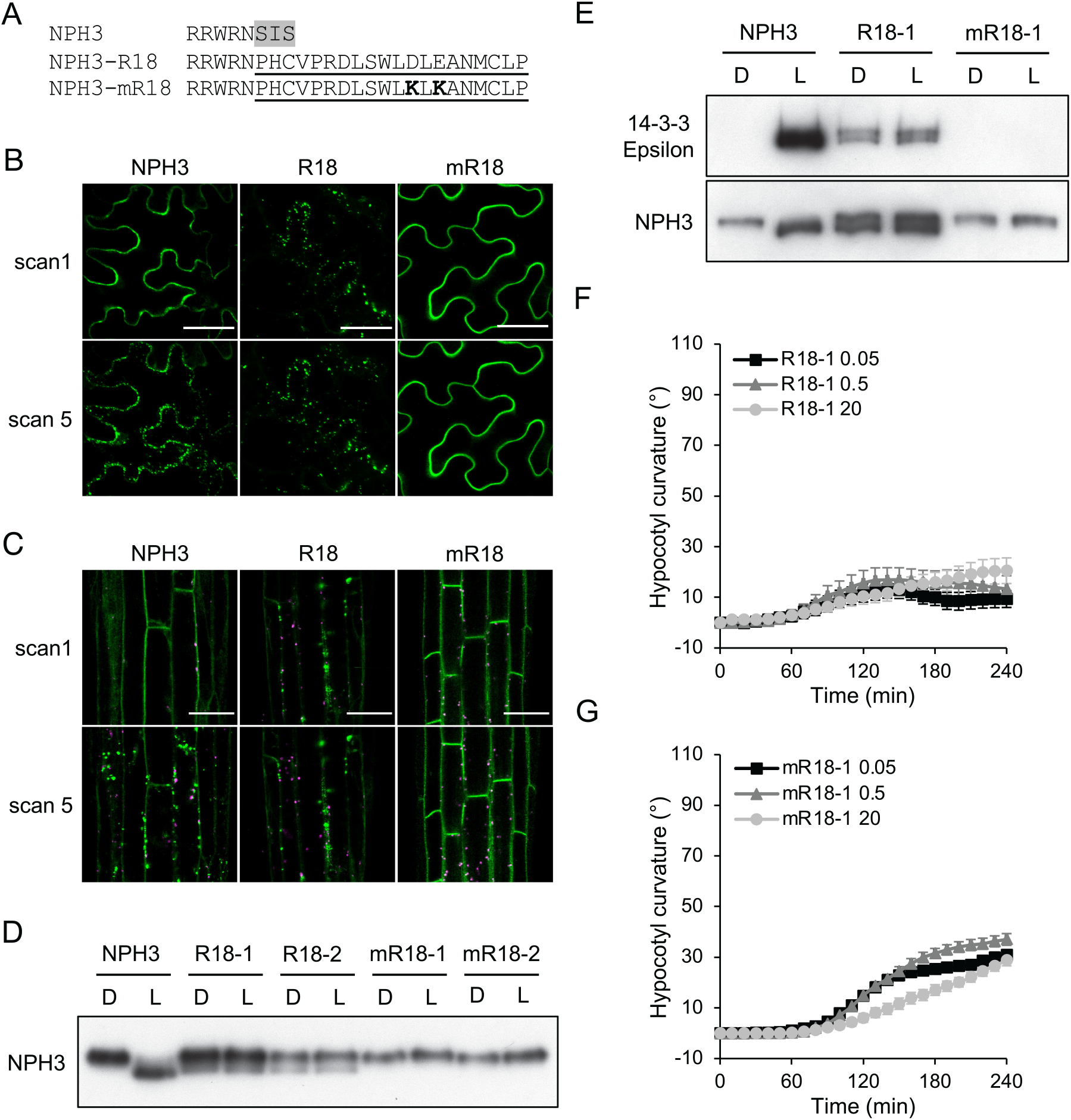
Analysis of a constitutive 14-3-3 binding NPH3 variant. (A) Amino acid sequence of the NPH3-R18 and mR18 constructs. Residues 744 – 746 of NPH3 (grey shaded) were replaced with the R18 peptide sequence (underlined). Two lysine residues (bold) were introduced into the mR18 sequence abolish 14-3-3 binding. Confocal images of GFP-NPH3 (NPH3), GFP-NPH3 containing the R18 peptide sequence (R18) or the mutated R18 peptide sequence (mR18) (B) transiently expressed in leaves of *N. benthamiana* plants, dark-adapted before confocal observation and (C) in hypocotyl cells of etiolated transgenic *Arabidopsis* seedlings. Images acquired immediately (scan 1) and after repeat scanning with the 488 nm laser (scan 5). GFP is shown in green and autofluorescence in magenta. Bar, 50 μm. (D) Immunoblot analysis of total protein extracts from etiolated seedlings expressing NPH3, R18 or mR18 maintained in darkness (D) or irradiated with 20 μmol m^−2^ s^−1^ blue light for 15 min (L). Protein extracts were probed with anti-NPH3 antibodies. (E) Far-western blot analysis of anti-GFP immunoprecipitations from etiolated seedlings expressing NPH3, R18 or mR18 maintained in darkness (D) or irradiated with 20 μmol m^−2^ s^−1^ of blue light for 15 min blue light (L). GST-tagged 14-3-3 isoform Epsilon was used as the probe. Blots were probed with anti-NPH3 antibody as loading control (bottom panel). Phototropism of etiolated seedlings expressing (F) R18 or (G) mR18 irradiated with 0.05 μmol m^−2^ s^−1^, 0.5 μmol m^−2^ s^−1^ or 20 μmol m^−2^ s^−1^ unilateral blue light. Hypocotyl curvatures were measured every 10 min for 4 h, and each value is the mean ± SE of 20 seedlings.

To confirm these results in stable transgenic lines, *Arabidopsis nph3* mutants were transformed with *NPH3::GFP-NPH3* containing the R18 or mR18 sequences. Confocal imaging of hypocotyl cells of etiolated seedlings revealed similar patterns of localisation observed in *N. benthamiana*, with GFP-NPH3-R18 forming aggregates in darkness, whereas GFP-NPH3-mR18 failed to relocalise following repeated laser scanning (Fig. 7C). Consistent with the subcellular localisation patterns, analysis of NPH3 dephosphorylation showed that lines expressing GFP-NPH3-mR18 display no change in electrophoretic mobility following blue light treatment, while a portion of GFP-NPH3-R18 exhibited enhanced electrophoretic mobility both in darkness and after irradiation (Fig. 7D). Far-western blotting was used to confirm the constitutive binding of recombinant 14-3-3 Epsilon to GFP-NPH3-R18 immunoprecipitated from seedlings maintained in darkness and following blue light irradiation, as well as the absence of 14-3-3 binding to GFP-NPH3-mR18 (Fig. 7E). Together these results show that engineered 14-3-3 binding, independent from phot1-medaited S744 phosphorylation, is partially sufficient to induce changes in NPH3 dephosphorylation and localisation status.

To assess functionality, phototropism was measured in GFP-NPH3-R18 and GFP-NPH3-mR18 expressing seedlings irradiated with 0.05 μmol m^−2^ s^−1^, 0.5 μmol m^−2^ s^−1^ or 20 μmol m^−2^ s^−1^ of unilateral blue light. Phototropic responsiveness was reduced under all fluence rates for GFP-NPH3-mR18 expressing seedlings (Fig. 7G, Fig. S3B), matching the phenotype of seedlings expressing GFP-NPH3 containing the S744A mutation (Fig. 6B - D). Phototropism was further reduced in seedlings expressing GFP-NPH3-R18 (Fig. 7F, Fig. S3A), which also displayed an increased variability in the direction of curvature (Fig. S3C) compared to the GFP-NPH3-mR18 lines. Therefore, while NPH3 mutants unable to bind 14-3-3 proteins have a reduced ability to reorientate growth towards a light source, constitutively 14-3-3 bound NPH3 mutants also display a diminished ability to sense the directionality of a light source.

## Discussion

In this study we used mass spectrometry to identify proteins co-immunoprecipitating with GFP-NPH3. This revealed 14-3-3 proteins as NPH3 interactors specifically following a blue-light treatment (Fig. 1). Using a chemical-genetic approach, we have found that NPH3 is phosphorylated by phot1 on the C-terminally positioned S744 in a light-dependent manner (Fig. 4A). Moreover, generation of anti-pS744 antibodies confirmed light-induced phosphorylation of S744 *in vivo* (Fig. 4C). Phototropins are members of the AGCVIII (protein kinase A, cyclic GMP-dependent protein kinase and protein kinase C) subfamily of protein kinases (Barbosa and Schwechheimer, 2014) and S744 is part of a PKA-like phosphorylation consensus sequence (RxS), as are the previously identified phot1-kinase substrates BLUS1 (Takemiya et al., 2013), CBC1 (Hiyama et al., 2017) and PKS4 (Schumacher et al., 2018); Fig. S4A).

Phot1-mediated phosphorylation of S744 is required to elicit the previously documented early cellular events associated with NPH3 activation such as dephosphorylation (Pedmale and Liscum, 2007) and subcellular relocalisation (Haga et al., 2015; Sullivan et al., 2019). This is consistent with previous observations of changes in NPH3 electrophoretic mobility correlating with the lifetime duration of phot1 activation *in planta* (Hart et al., 2019) and occurring locally only in cells/tissues where both proteins are present (Sullivan et al., 2016). Furthermore, a constitutively active phot1-variant can induce NPH3 dephosphorylation in darkness (Kimura et al., 2020). The phosphorylation status of residues S213, S223, S233 and S237 contribute to reducing the electrophoretic mobility of NPH3 in darkness (Tsuchida-Mayama et al., 2008), however other unidentified sites are also involved (Fig. 2C; (Haga et al., 2015). The kinase(s) and phosphatase(s) regulating the phosphorylation status of these sites is currently unknown, as is their role in regulating NPH3 signalling. However, mutation of S213, S223, S233 and S237 to alanine, or deletion of amino acid residues S213-S239, did not impact their ability to restore phototropism in *nph3* mutant seedlings (Tsuchida-Mayama et al., 2008), or form aggregates when transiently expressed in *N. benthamiana* (Fig. 2B).

Phosphorylation of S744 creates a 14-3-3 binding site (Fig. 3C) which conforms to the C-terminal mode III 14-3-3 binding motif pS/pTX_1-2_-COOH (Camoni et al., 2018). We created a translational fusion between NPH3 and the synthetic R18 peptide to study the role of 14-3-3 binding in the absence of phot1 phosphorylation (Fig. 7E). 14-3-3 binding alone was able to induce NPH3 relocalisation into aggregates (Fig. 7B, Fig. 7C) and partially reduce the electrophoretic mobility of NPH3 (Fig. 7D), in the absence or presence of light. Light-dependent 14-3-3 binding has also been shown for phot1; non-epsilon 14-3-3s bind to 3 phosphorylation sites located between the LOV1 and LOV2 photosensory domains (Sullivan et al., 2009), but the functional relevance of this interaction is unknown as mutation of 2 of the phosphorylation sites did not impair functionality (Inoue et al., 2008a). In contrast, no isoform specificity was observed for 14-3-3 binding to NPH3, with both epsilon and non-epsilon isoforms shown to interact (Fig. 1). Functional redundancy between 14-3-3 isoforms means loss-of-function mutants often show few, if any, phenotypes, with even quadruple non-epsilon 14-3-3 mutants displaying mild growth phenotypes under non-stress growth conditions (van Kleeff et al., 2014), with no obvious differences in phototropism or NPH3 dephosphorylation kinetics observed compared to WT seedlings (Fig. S5). However, conditional RNA interference (RNAi) lines targeting three 14-3-3 epsilon members (epsilon, mu and omicron) displayed several auxin-related phenotypes, including reduced hypocotyl elongation and defects in root and hypocotyl gravitropism, due to altered polarity of the PIN-FORMED (PIN) auxin transporters as a consequence of 14-3-3 regulation of cellular trafficking (Keicher et al., 2017). NPH3 is also reported to be required for phot1-driven changes in PIN2 trafficking during negative phototropic bending of roots (Wan et al., 2012). However, the role of asymmetric auxin distribution in root phototropism has recently been questioned (Kimura et al., 2018).

The biochemical basis underpinning phototropism is the formation of a gradient of phot1 activation across the stem (Salomon et al., 1997), which results in an asymmetric accumulation of auxin on the shaded side through an unidentified mechanism (Fankhauser and Christie, 2015). We previously demonstrated that a gradient of GFP-NPH3 relocalisation occurs across the hypocotyl of *Arabidopsis* seedlings during unilateral irradiation with blue light (Sullivan et al., 2019). Here we report that seedlings expressing mutants of GFP-NPH3 unable to form such a gradient, either through mutation of the phosphorylation site required for 14-3-3 binding (S744) or due to constitutive 14-3-3 binding via the R18 peptide, have a severely compromised phototropic response. Thus, phototropic curvature likely involves signalling outputs mediated by a gradient in NPH3 localisation across the stem. Our current findings are therefore consistent with 14-3-3 proteins being instrumental components regulating auxin-dependent growth (Keicher et al., 2017).

The phot1 phosphorylation consensus sequence of NPH3 is also conserved in several other NRL proteins including RPT2, NCH1 and members of the NAKED PINS IN YUCCA (NPY) clade (Fig. S4B). Notably, RPT2 was identified in immunoprecipitants of seedlings expressing 14-3-3 epsilon-GFP (Keicher et al., 2017). We could also detect phosphorylation of RPT2 on the corresponding serine residue (S591) when co-expressed with phot1^GK^ in *in vitro* kinase assays (Fig. S4C). It is therefore possible the residual functionality seen in GFP-NPH3 S744A seedlings (Fig. 6) arises from co-action with other NRL family members. However dynamic relocalisation in response to blue light has not been reported for RPT2 (Haga et al., 2015; Kimura et al., 2020) or NCH1 (Suetsugu et al., 2016), hence the consequences of phosphorylation and 14-3-3 binding must differ for specific NRL family members. The NPY clade of NRL proteins function redundantly to mediate organogenesis and root gravitropism (Furutani et al., 2007; Furutani et al., 2011; Li et al., 2011). These responses are independent of phototropin signalling but involve related AGCVIII kinases PINOID (PID) and its close homologues WAG1 and WAG2, and the D6 PROTEIN KINASE (D6PK) family (Glanc et al., 2021). PID/WAGs and D6PKs phosphorylate PIN transporters on RxS phosphorylation site motifs (Barbosa and Schwechheimer, 2014) and physically interact with NPY proteins (Glanc et al., 2021). Furthermore, aggregate formation is not limited to NPH3 and has been documented for NPY1 when expressed in *Arabidopsis* protoplasts (Furutani et al., 2007). Therefore, phosphorylation and concomitant 14-3-3 binding to the C-terminus may represent a conserved mechanism of regulation for NRL proteins.

Determining the biochemical function of NPH3 is now required to understand how phots signal via NRL proteins to coordinate different light-capturing processes in plants that will ultimately offer new opportunities to manipulate plant growth through alterations in photosynthetic capacity.

## METHODS

### Plant Material and growth

Wild-type *Arabidopsis* (*gl-1*, ecotype Columbia), *nph3-6* (Motchoulski and Liscum, 1999), 14-3-3 quadruple mutants (van Kleeff et al., 2014) and the GFP-NPH3 transgenic line (Sullivan et al., 2019) were previously described. Unless otherwise stated, seeds were sown on soil or surface sterilised and planted on half-strength Murashige and Skoog (MS) medium with 0.8% agar (w/v) and stratified at 4°C for 2 - 5 d. Seeds on soil were transferred to a controlled environment room (Fitotron, Weiss Technik) with LED illumination (C65NS12, Valoya) under 16 h 22 °C/ 8 h 18 °C light: dark cycles and 80 μmol m^−2^ s^−1^ white light. Seeds on MS medium were exposed to 80 μmol m^−2^ s^−1^ white light for 6 to 8 h to induce germination and grown vertically in darkness for 3 d. For blue light treatment, white light was filtered through Moonlight Blue filter No. 183 (Lee Filters). Fluence rates for all light sources were measured with an Li-250A and quantum sensor (LI-COR).

### Transient Expression in *Nicotiana benthamiana*

To create transformation vectors encoding *NPH3* with multiple serine and threonine residues mutated to alanine, fragments of *NPH3* were synthesised (ThermoFisher Scientific) encoding the 13 alanine substitutions for *NPH3-M1*, 8 substitutions for *NPH3-M2* and 15 substitutions for *NPH3-M3*. The synthesised fragments were introduced into *NPH3::GFP-NPH3* using *KpnI* and *MluI* restriction sites for *GFP-NPH3-M1, MluI* and *PstI* restriction sites for *GFP-NPH3-M2* and *PstI* and *BamHI* restriction sites for *GFP-NPH3-M3. Agrobacterium-mediated* transient expression in *Nicotiana benthamiana* was performed as previously described (Kaiserli et al., 2009). *Agrobacterium tumefaciens* strain GV3101, transformed with the plasmid of interest, was resuspended in infiltration buffer (10 mM MgCl2, 10 mM MES-KOH [pH 5.6], and 200 mM acetosyringone) at an OD_600_ of 0.4 and syringe-infiltrated into leaves of 3 to 4-week-old *N. benthamiana* plants. Plants were dark-adapted for 16 h before 1 cm leaf discs for confocal observation or protein extraction were taken 2 d post-infiltration. For blue-light irradiation, leaf discs were placed abaxial-side upwards on the surface of MS medium agar plates for the duration of the treatment.

### Transformation of Arabidopsis

Amino acid substitutions of S744 and/or S746 were introduced into the pUC-SP vector containing the *NPH3* coding sequence by site-directed mutagenesis and verified by DNA sequencing. The coding sequence of *NPH3* in the *NPH3::GFP-NPH3* pEZR(K)-LC binary vector (Sullivan et al., 2019) was replaced by the coding sequences containing the phosphosite mutations using Gibson Assembly (New England Biolabs). To create transformation vectors *NPH3::GFP-NPH3-R18* and *NPH3::GFP-NPH3-mR18*, a fragment encoding amino acid residues 419 – 743 was PCR amplified from *NPH3* pUC-SP with primers containing the R18 or mR18 coding sequence and inserted into *NPH3::GFP-NPH3* using *MluI* and *BamHI* restriction sites. The *nph3-6* mutant was transformed with *Agrobacterium tumefaciens* strain GV3101 as previously described (Davis et al., 2009). Based on the segregation of kanamycin resistance independent homozygous T3 lines, or for GFP-NPH3-mR18 transgenics single-insertion T2 lines, were selected for analysis.

### Phototropism

Phototropism was performed using free-standing etiolated seedlings grown on a layer of silicon dioxide (Honeywell, Fluka), watered with quarter-strength MS medium, as previously described (Sullivan et al., 2016). Images were recorded every 10 min for 4 h with a Retiga 6000 CCD camera (QImaging) connected to a personal computer running QCapture Pro 7 software (QImaging) with supplemental infrared illumination. Hypocotyl curvature was measured using Fiji software (Schindelin et al., 2012). Circular histograms were produced using Oriana software (Kovach Computing Services).

### Leaf positioning

Seedlings were grown on soil for 9 d under 80 μmol m^−2^ s^−1^ white light before transfer to 10 μmol m^−2^ s^−1^ white light for 4 d. One cotyledon was removed, seedlings were placed flat on an agar plate, and plates were placed on a white light transilluminator and photographed. Petiole angles from the horizontal were measured using Fiji software.

### Confocal Microscopy

Localization of GFP-tagged NPH3 was visualized with a Leica SP8 laser scanning confocal microscope using HC PL APO 20x/0.75 or 40×/1.30 objective. The 488-nm excitation line was used, and GFP fluorescence was collected between 500 and 530 nm. Images were acquired at 1,024-1,024-pixel resolution with a line average of two. Z-stacks were acquired at 5 min intervals, with darkness between each scan. Maximum projection images were constructed from z-stacks using Fiji software.

### Immunoblot Analysis

Total proteins were extracted by grinding 50 3-d-old etiolated *Arabidopsis* seedlings or 1 cm *Agrobacterium*-infiltrated *N. benthamiana* leaf discs in 100 μl 2 X SDS sample buffer under red safe light illumination, clarified by centrifugation at 13 000 g for 5 min, boiled for 4 min and subjected to SDS-PAGE. Proteins were transferred onto a polyvinylidene fluoride membrane (PVDF; Bio-Rad) with a Trans-Blot Turbo Transfer System (Bio-Rad) and detected with anti-GST monoclonal antibody (Merck), anti-GFP-HFP monoclonal antibody (Miltenyi Biotech), anti-thiophosphoester monoclonal antibody (clone 51-8, Abcam), anti-UGPase antibody (AgriSera), anti-phot1 polyclonal antibodies (Cho et al., 2007), anti-NPH3 purified polyclonal antibodies raised against peptides IPNRKTLIEATPQSF and GVDHPPPRKPRRWRN (Eurogentec) and polyclonal antibodies raised against phosphorylated S744 of NPH3 using peptide KPRRWRNpSIS (where pS represents phosphorylated serine) as antigen (Eurogentec). Blots were developed with horseradish peroxidase (HRP)-linked secondary antibodies (Promega) and Immobilon Western Chemiluminescent HRP Substrate (Merck).

### Far-Western Blot Analysis

*Arabidopsis* 14-3-3 isoforms Epsilon and Lambda were expressed using the pGEX-4T1 vector (Merck), as a translational fusion with glutathione-S-transferase (GST) and purified with GST-Bind resin (Merck), as previously described (Sullivan et al., 2009). Total protein extracts were prepared from 3-d-old etiolated seedlings maintained in darkness, or following blue-light irradiation, under a dim red safe light. Seedlings were ground in a mortar and pestle in GTEN buffer (10% [v/v] glycerol, 25 mM Tris-HCl [pH 7.5], 1 mM EDTA, 150 mM NaCl) supplemented with 0.5% SDS, 10 mM DTT, 1 mM phenylmethylsulfonyl fluoride (PMSF) and a protease inhibitor mixture (Complete EDTA-free; Merck) on ice and clarified by centrifugation at 10 000 g, 4 °C for 10 min. Immunoprecipitations were preformed using GFP-Trap Agarose beads (Chromotek), eluted by boiling in 2 X SDS sample buffer, separated by SDS-PAGE and transferred to PVDF membrane. PVDF membranes were incubated with purified GST-14-3-3 proteins or GST alone in far-western buffer (20 mM HEPES-KOH [pH 7.7], 75 mM KCl, 0.1 mM EDTA, 1 mM DTT, 2% milk, 0.04% Tween-20) at a final concentration of 1 μM. 14-3-3 binding was detected using anti-GST monoclonal antibody (Merck).

### Immunoprecipitation

Total protein extracts were prepared from 3-d-old etiolated WT seedlings or seedlings expressing GFP-NPH3 maintained in darkness (Dark) or irradiated with 20 μmol m^−2^ s^−1^ of blue light for 15 min. Seedlings were ground in a mortar and pestle in IP buffer (50 mM Tris-HCl [pH 7.5], 150 mM NaCl, 1 mM EDTA, 1% Triton X-100, 1 mM PMSF) supplemented with protease inhibitor mixture (Complete EDTA-free; Merck) and half-strength phosphatase inhibitor cocktail 2 and 3 (Merck). Samples were clarified twice by centrifugation at 14 000 g, 4 °C for 10 min. Immunoprecipitations were preformed using the μMACS GFP isolation kit (Miltenyi Biotech), eluted with 0.1 M Triethylamine pH 11.8/0.1% Triton X-100 and neutralized with 1 M MES pH 3. Proteins were identified by liquid chromatography–tandem mass spectrometry using the Fingerprints Proteomics Facility (University of Dundee). Only proteins identified in 2 biological replicates from at least 2 peptides were retained for analysis. Proteins identified in immunoprecipitations from WT seedlings were considered contaminants. Protein intensities were converted to relative abundance of the bait protein (GFP-NPH3), which was set to 100 in each sample. Proteins showing at least a two-fold change in relative abundance following blue-light irradiation were identified (Table S1).

### *In vitro* kinase assay

The coding sequence of *NPH3* in *NPH3::GFP-NPH3* and *NPH3::GFP-NPH3 S744A* was amplified and inserted into the pSP64 poly(A) vector (Promega), together with a N-terminal Haemagglutinin (HA) tag, using Gibson Assembly (New England Biolabs). The *RPT2* coding sequence was amplified and inserted into the pSP64 poly(A) vector, together with a N-terminal GST tag, using Gibson Assembly (New England Biolabs). The RPT2 S591A substitution was introduced by site-directed mutagenesis. *In vitro* kinase assays were performed by co-expressing the substrate together with a gate-keeper engineered phot1 (phot1^GK^) using the TnT^®^ SP6 High-Yield Wheat Germ Protein Expression System (Promega) in the presence of 10 μM FMN, as previously described (Schnabel et al., 2018). Thiophosphorylation reactions were performed in the presence of 500 μM N^6^-benzyl-ATPγS (Jena Bioscience), in phosphorylation buffer contained 37.5 mM Tris-HCl pH 7.5, 5.3 mM MgSO4, 150 mM NaCl and 1 mM EGTA. Samples were either mock irradiated or treated for 20 s with white light at a total fluence of 60,000 μmol m^−2^. Reactions were performed for 5 min and stopped by addition of EDTA (pH 8.0) to a final concentration of 20 mM. Thiophosphorylated molecules were alkylated with 2.5 mM p-nitrobenzyl mesylate (PNBM, Abcam) for 2 hours. HA-tagged NPH3 was immunoprecipitated using Pierce™ Anti-HA Magnetic Beads (Thermo Fisher Scientific). Thiophosphorylation was visualised by immunoblotting with anti-thiophosphoester monoclonal antibody (clone 51-8, Abcam).

## Acknowledgements

This work was supported by funding from the UK Biotechnology and Biological Sciences Research Council (BB/M002128/1, BB/R001499/1 to J.M.C) and the Grant-in-Aid for Scientific Research Grant from the Japan Society for the Promotion of Science (15KK0254 to N.S.). We are grateful to Albertus H. de Boer for providing 14-3-3 quadruple mutant seed. We are indebted to Claudia Oecking for sharing data and helpful discussions.

## Author Contributions

S.S, N.S. and J.M.C designed research; S.S., T.W., L.H., D.P. and M.L. performed research; S.S, N.S. and J.M.C analysed data; S.S and J.M.C wrote the manuscript. All authors commented on the manuscript.

---

**Table S1.** GFP-NPH3 interacting proteins identified by mass spectrometry analysis of anti-GFP immunoprecipitations.

**Fig. S1.**
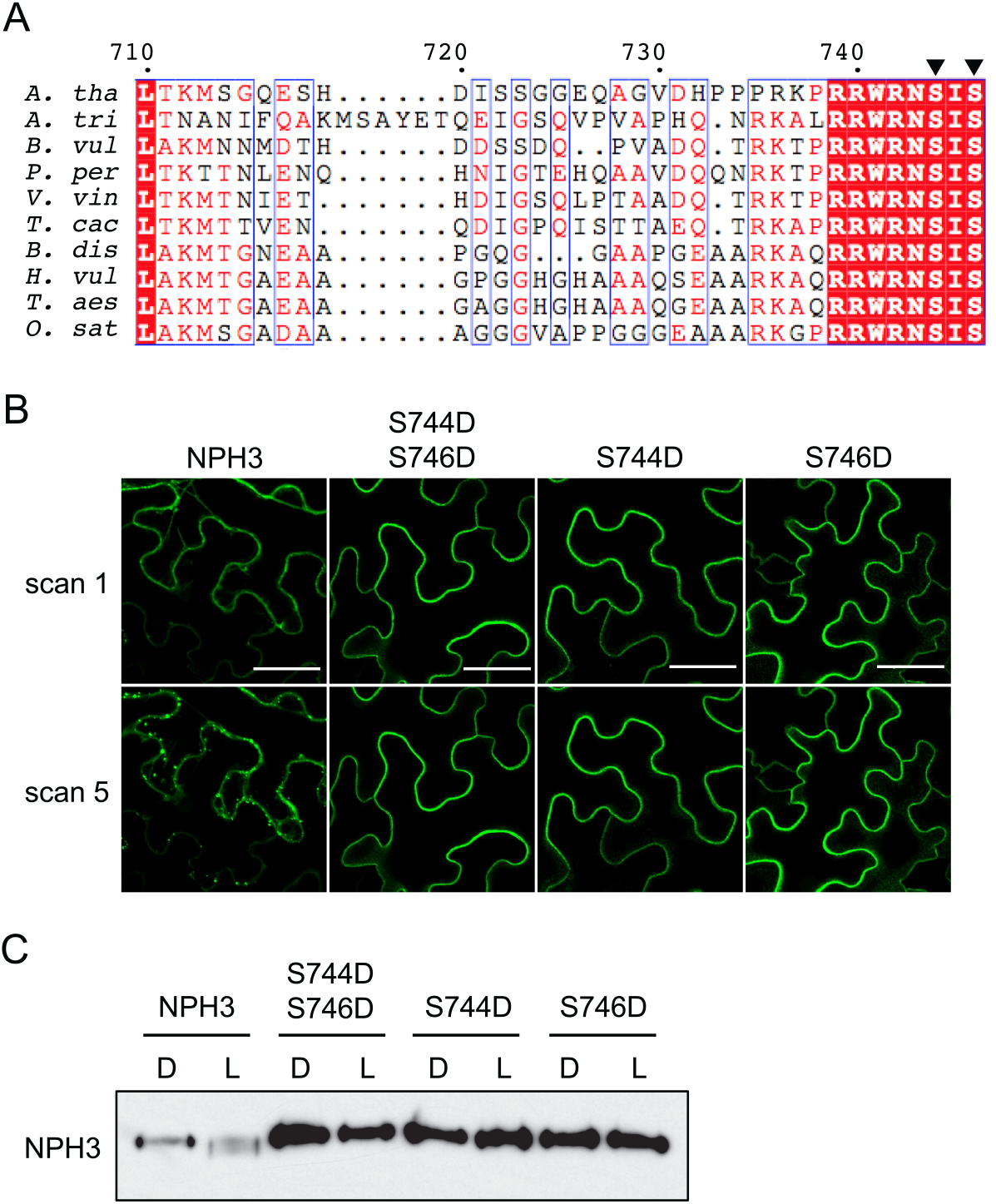
Mutational analysis of NPH3 phosphorylation sites. (A) Amino acid alignment of the C-terminus of NPH3 from *Arabidopsis thaliana (A. tha), Amborella trichopoda (A. tri), Beta vulgaris (B. vul), Prunus persica (P. per), Vitis vinifera (V. vin), Theobroma cacao (T. cac), Brachypodium distachyon (B. dis), Hordeum vulgare (H. vul) Triticum aestivum (T. aes) and Oryza sativa (O. sat)*. The two conserved serine residues (*A. tha*, S744 and S746) are indicated by arrow heads. (B) Confocal images of GFP-NPH3 (NPH3) and phosphorylation site mutants S744D S746D, S744D and S746D transiently expressed in leaves of *N. benthamiana*. Plants were dark-adapted before confocal observation and images acquired immediately (scan 1) and after repeat scanning with the 488 nm laser (scan 5). Bar, 50 μm. (C) Immunoblot analysis of protein extracts from leaves of *N. benthamiana* transiently expressing GFP-NPH3 (NPH3) and phosphorylation site mutants S744D S746D, S744D and S746D. Plants were dark-adapted and maintained in darkness (D) or irradiated with 20 μmol m^−2^ s^−1^ of blue light for 15 min (L). Protein extracts were probed with anti-GFP antibodies.

**Fig. S2.**
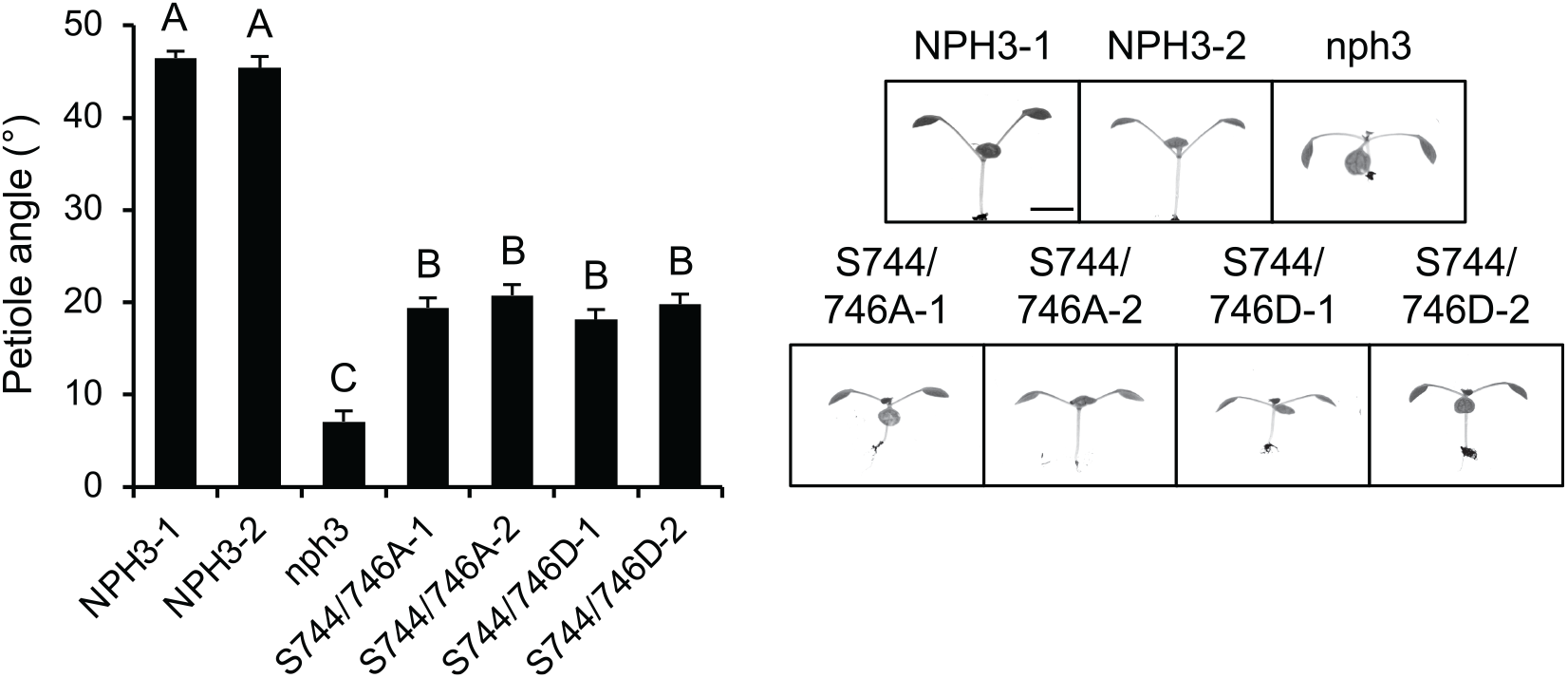
Phot1 phosphorylation of NPH3 promotes functionality. (A) Petiole positioning of *nph3* mutant and seedlings expressing GFP-NPH3 (NPH3) or phosphorylation site mutants S744A S746A and S744D S746D. Plants were grown under 80 μmol m^−2^ s^−1^ white light for 9 d before transfer to 10 μmol m^−2^ s^−1^ white light for 5 d. Petiole angle from the horizontal was measured for the first true leaves, each value is the mean ± SE of 20 seedlings. Means that do not share a letter are significantly different (P < 0.01, one-way ANOVA with Tukey HSD post-test). Representative images for each genotype are shown on the right. Bar, 5 mm.

**Fig. S3.**
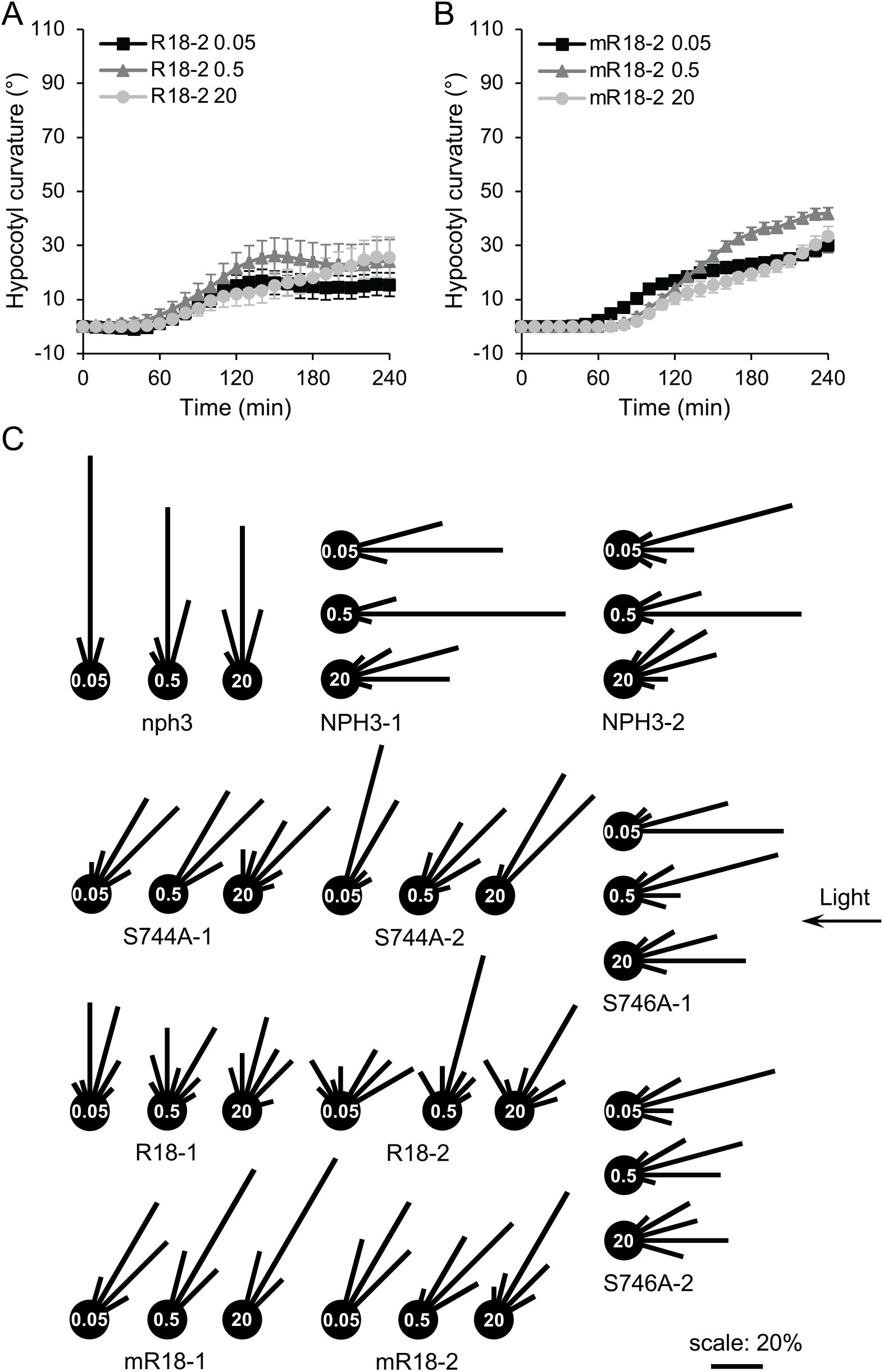
Analysis of a constitutive 14-3-3 binding NPH3 variant. Phototropism of etiolated seedlings expressing (A) R18 or (B) mR18 irradiated with 0.05 μmol m^−2^ s^−1^, 0.5 μmol m^−2^ s^−1^ or 20 μmol m^−2^ s^−1^ unilateral blue light. Hypocotyl curvatures were measured every 10 min for 4 h, and each value is the mean ± SE of 18-20 seedlings. (C) Circular histograms depicting hypocotyl orientation after 240 min of irradiation with 0.05 μmol m^−2^ s^−1^, 0.5 μmol m^−2^ s^−1^ or 20 μmol m^−2^ s^−1^ unilateral blue light. Angles were grouped into 15° classes and expressed as percentages of the number of seedlings.

**Fig. S4.**
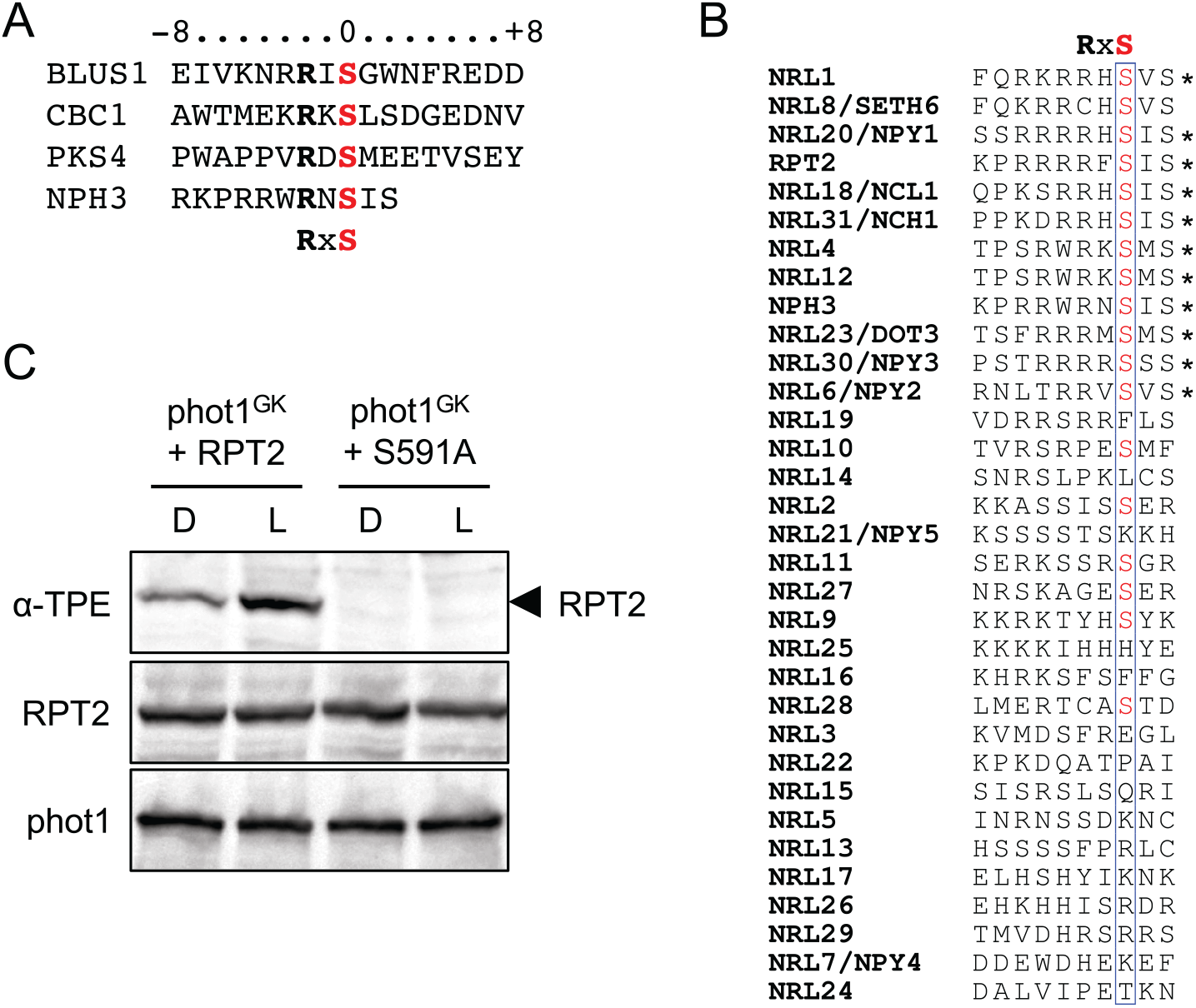
Conservation of the phot1 phosphorylation sequence of NPH3. (A) Amino acid sequence alignment of the phototropin1 substrate phosphorylation sites in BLUS1 (Takemiya et al., 2013), CBC1 (Hiyama et al., 2017), PKS4 (Schumacher et al., 2018) and NPH3. The amino acid residues are numbered relative to the phosphorylated serine residue and the PKA-like phosphorylation motif is indicated below. (B) Amino acid alignment of the last 10 residues of the *Arabidopsis* NRL protein family. The position of the RxS phosphorylation motif is indicated above and sequences containing a RxS motif denoted with an asterisk. (C) Thiophosphorylation analysis of *in vitro* kinase assays containing gatekeeper engineered phot1 (phot1^GK^) and RPT2 or RPT2-S591A. Reactions were performed in the absence (D) or presence of 20 s of white light (L), and thiophosphorylation was detected using anti-thiophosphoester antibody (α-TPE). Blots were probed with anti-GST antibody to detect GST-RPT2 and phot1 ^GK_^ GST.

**Fig. S5.**
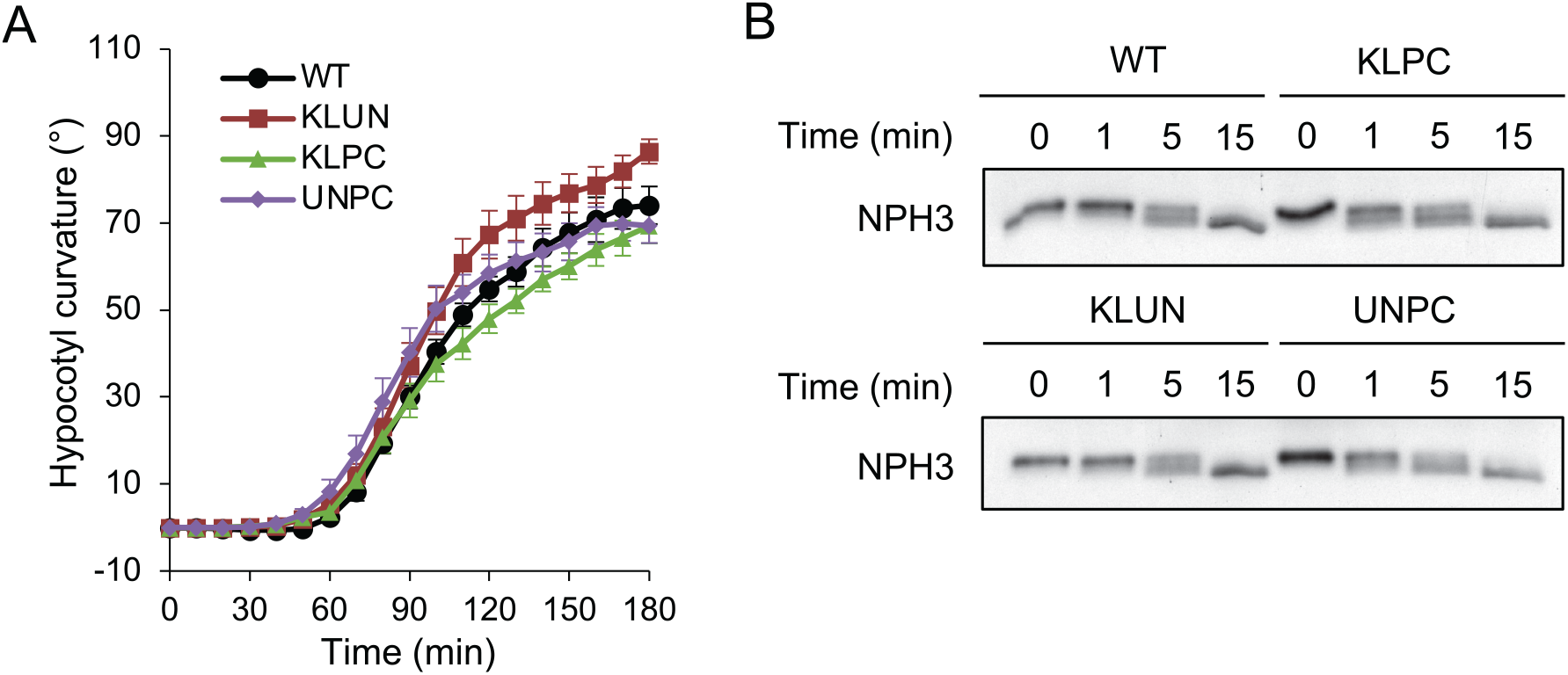
Analysis of quadruple 14-3-3 mutants. (A) Phototropism of etiolated wild-type (WT) seedlings or *kappa lambda phi chi* (KLPC), *kappa lambda upsilon nu* (KLUN) and *upsilon nu phi chi* (UNPC) quadruple mutant seedlings irradiated with 0.5 μmol m^−2^ s^−1^ unilateral blue light. Hypocotyl curvatures were measured every 10 min for 3 h, and each value is the mean ± SE of 10 seedlings. (B) Immunoblot analysis of total protein extracts from etiolated WT or KLPC, KLPC and UNPC quadruple mutant seedlings irradiated with 15 μmol m^−2^ s^−1^ of blue light for the time indicated. Blots were probed with anti-NPH3 antibodies.

